# lmeEEG: Mass linear mixed-effects modeling of EEG data with crossed random effects

**DOI:** 10.1101/2023.01.18.524560

**Authors:** Antonino Visalli, Maria Montefinese, Giada Viviani, Livio Finos, Antonino Vallesi, Ettore Ambrosini

**Affiliations:** IRCCS San Camillo Hospital, Venice, Italy; Department of Developmental and Social Psychology, University of Padova, Padova, Italy; Department of Neuroscience, University of Padova, Padova, Italy; Padova Neuroscience Center, University of Padova, Padova, Italy; Department of Statistical Sciences, University of Padova, Padova, Italy; Department of General Psychology, University of Padova, Padova, Italy

**Keywords:** EEG, linear mixed-effects models, TFCE, mass-univariate testing, crossed random effects, Psycholinguistics

## Abstract

**Background:** Mixed-effects models are the current standard for the analysis of behavioral studies in psycholinguistics and related fields, given their ability to simultaneously model crossed random effects for subjects and items. However, they are hardly applied in neuroimaging and psychophysiology, where the use of mass univariate analyses in combination with permutation testing would be too computationally demanding to be practicable with mixed models.

**New method:** Here, we propose and validate an analytical strategy that enables the use of linear mixed models (LMM) with crossed random intercepts in mass univariate analyses of EEG data (lmeEEG). It avoids the unfeasible computational costs that would arise from massive permutation testing with LMM using a simple solution: removing random-effects contributions from EEG data and performing mass univariate linear analysis and permutations on the obtained marginal EEG.

**Results:** lmeEEG showed excellent performance properties in terms of power and false positive rate.

**Comparison with existing methods:** lmeEEG overcomes the computational costs of standard available approaches (our method was indeed more than 300 times faster).

**Conclusions:** lmeEEG allows researchers to use mixed models with EEG mass univariate analyses. Thanks to the possibility offered by the method described here, we anticipate that LMM will become increasingly important in neuroscience. Data and codes are available at osf.io/kw87a. The codes and a tutorial are also available at github.com/antovis86/lmeEEG.

## 1. Introduction

Mixed-effects models are crucial to appropriately analyze data from experimental designs including both subjects and items as crossed random effects (Baayen et al., 2008; DeBruine & Barr, 2021). However, their use is limited in neuroimaging and electrophysiological data analyses, also due to computational time constraints. Here, we introduce an analytical strategy for performing mass univariate linear mixed-effects model analyses of EEG data (lmeEEG), such as event-related brain potentials.

Consider, as an example of a design with crossed random effects, a psycholinguistic study in which participants are asked to judge a linguistic property of a set of words. When analyzing psychological experiments, researchers take into account inter-individual variability (or random error) to draw general conclusions that are valid beyond the selected sample (this aspect is implicit in the majority of repeated-measures statistical models). It is noteworthy that linguistic stimuli are also sampled from the population of all words. As participants are treated as random variables to generalize results to their population, the same logic applies to items (Barr, 2017; Clark, 1973). Indeed, researchers are usually not interested in experimental effects that are valid only for the selected set of stimuli used in the specific study (words in this example). Hence, inter-item variability must be considered to generalize results to the population of words from which the experimental items are drawn. Although modeling by-item variability has mainly attracted the attention of psycholinguistics, doing this is mandatory whenever a researcher analyzes data from an experimental study (e.g., memory or emotion studies) in which stimuli are drawn from a larger population in any field (Barr, 2017).

Given their ability to simultaneously model crossed random effects, mixed models are the current standard for behavioral studies in psycholinguistics and related fields (DeBruine & Barr, 2021), but they are less common in neuroimaging and psychophysiology. Focusing on electroencephalography (EEG) or magnetoencephalography (MEG), in recent years the field has moved from traditional approaches that analyze selected channels and timepoints to mass univariate approaches, in which the whole Channel(Sensor) × Timepoint data space is tested (Groppe et al., 2011; Woolrich et al., 2009). However, the use of mass univariate analyses in combination with resampling methods (i.e., permutation testing and bootstrapping) to control for the Family-Wise Error Rate (FWER) (Pernet et al., 2015) is overly computationally demanding to be practicable with mixed models (Fields & Kuperberg, 2020; Nielson & Sederberg, 2017). Indeed, linear mixed models (LMM) are estimated using (restricted) maximum likelihood ((RE)ML) methods, which require too much time to be performed millions of times. This is probably one of the reasons why the available toolboxes for mass univariate analysis of M-EEG data (e.g., LIMO EEG: Pernet et al., 2011; SPM: Kiebel & Friston, 2004; Unfold: Ehinger & Dimigen, 2019; Kherad-Pajouh & Renaud, 2015; Frossard, 2019; Frossard & Renaud, 2021) include only statistical tests that can be reconducted to linear models (LM), which rely on ordinary least squares (OLS) solutions (considerably faster than (RE)ML estimations). These toolboxes perform random coefficient analysis (also called two-step linear regression, multilevel linear model, or hierarchical general linear model; Lorch & Myers, 1990), in which fixed effects coefficients are first estimated within each participant and then tested at the group level. Although these models deal with inter-individual variability, they cannot simultaneously model crossed random effects. It follows that they are not appropriate for the analysis of experimental designs in which stimuli are a sample drawn from a larger population of items (Bürki et al., 2018).

Here, we propose a solution to the unfeasible computational costs derived from the use of permutation methods with LMM. Unlike other approaches that reduce the dimensionality of the analyzed EEG datasets before performing LMM (Nielson & Sederberg, 2017), thus preventing the possibility to fully exploit the entire spatio-temporal information in EEG data, our approach (lmeEEG) enables the use of mixed models with mass univariate analyses. lmeEEG complements other mass univariate modeling techniques by providing a method for analyzing experimental designs with crossed random effects. In the following, we first describe our method in detail. Secondly, we present a validation of lmeEEG using a simulated experiment. Lastly, we present its application to a real EEG dataset.

## 2. Description of lmeEEG

To introduce lmeEEG, we will describe its application to a simplified experiment. Three participants *s*1, *s*2, and *s*3 perform a semantic decision task (i.e., judging whether a word is abstract or concrete) on four words that belong to the experimental conditions concrete (w*1* and w*2*) and abstract (w*3* and w*4)*. Analyses are performed on EEG data collected from 19 channels at 110 timepoints time-locked to word onset.

lmeEEG consists of the following steps (Fig. 1):

**Figure 1.**
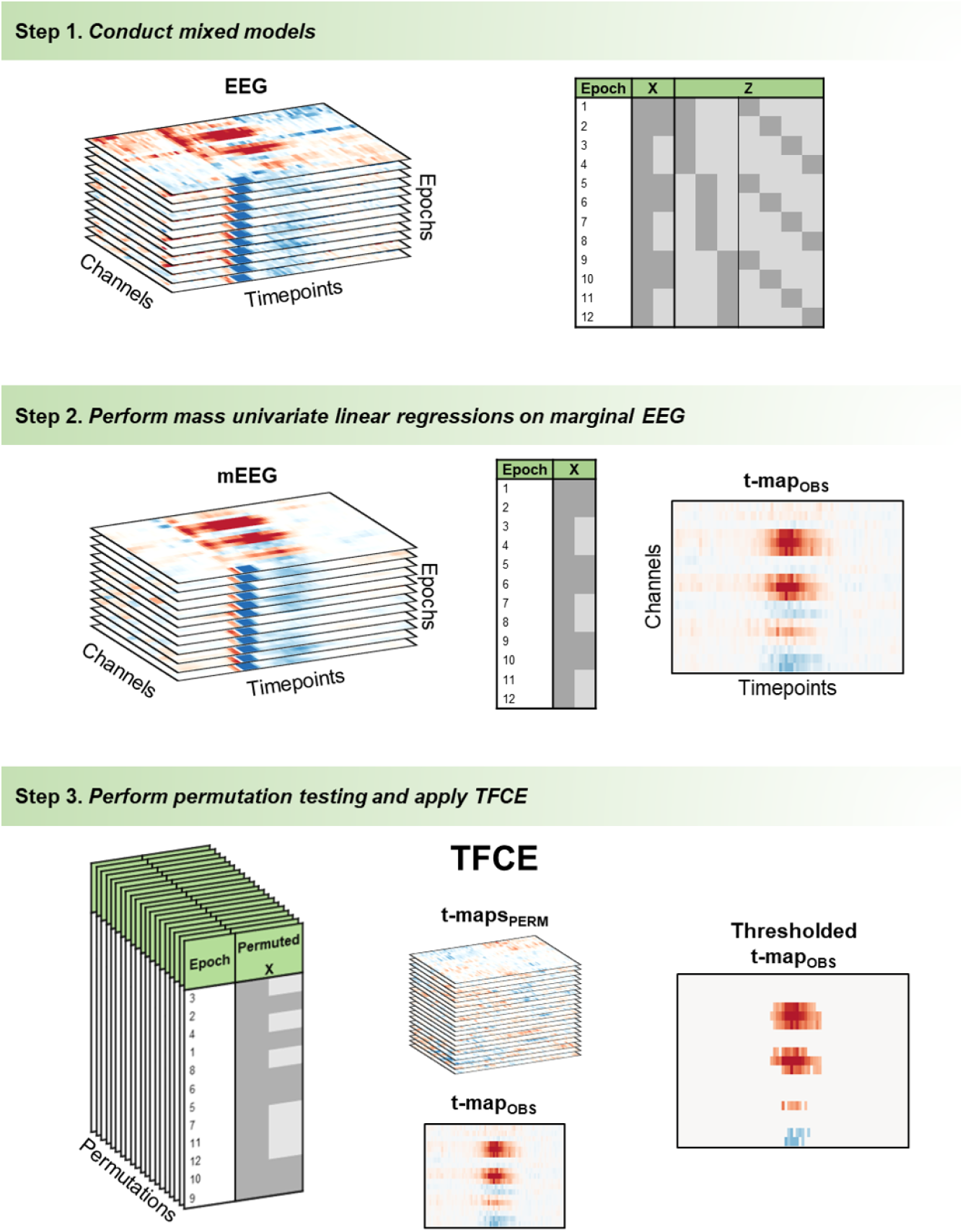
Illustration of the analytical steps in lmeEEG. In step 1 (top), a linear mixed model (LMM) is massively conducted at each channel and timepoint combination on epoched EEG data vectors comprising all trials from all subjects. The LMM design matrix is composed of a fixed-effects part (X) and a random-effects part (Z). In step 2 (middle), marginal EEG data (mEEG) vectors for each channel/timepoint pair are obtained by removing random effects contributions estimated in step 1. Mass univariate linear regression models (LM) - composed of only X - are conducted on mEEG and a map of *t* values (*t*-map_OBS_) is obtained for each predictor of interest (one predictor in the example). In step 3 (bottom), X is permutated and used for the mass LM. Then, threshold-free cluster enhancement (TFCE) is applied on the *t*-maps obtained from each permutation (*t*-maps_PERM_) and the maxTFCE values of each permutation used to build an empirical distribution of the maxTFCE values under H_0_. The empirical distribution of the maxTFCE values is used to assess the statistical significance of the TFCE values of *t*-map_OBS_.

### 1. Conduct mixed models on each channel/timepoint combination

For each EEG channel *ch* and event-locked timepoint *t*, a linear mixed-effects model (LMM) is conducted on trial-wise EEG responses concatenated across participants (EEG_*ch,t*_ in Eq.1). The LMM can be summarized as follows:

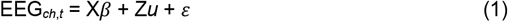

In (1), X represents the fixed-effects design matrix, which includes in the present example a column of ones for the intercept and a column representing the experimental factor contrast. The X matrix is multiplied by the population coefficients β, here consisting of β_0_ for the intercept and β_1_ for the contrast of abstract words compared to concrete words. Continuing with (1), Z represents the random-effects design matrix. In the proposed example, Z is composed of two grouping variables, a three-column variable for participants and a four-column variable for words, which are multiplied by the coefficients *u*, which indicate the value that must be added to the population intercept for each participant and each item. Finally, ε represents the residual. A remark needs to be made on the specification of the random-effects structure. In the example, we used a minimal structure, allowing only intercepts to vary across participants and words. Different approaches have been proposed for the random-effects specifications (Barr et al., 2013; Bates, Kliegl, et al., 2015; Matuschek et al., 2017), but it is important here to note that it is hard to assess and manage convergence and singularity issues with massive testing; and random-effects parameters are more difficult to estimate and their number rapidly increases as model complexity slightly increases, thus leading to important computational costs. In any case, here, we limit the validation of our method to models without random slopes.

### 2. Perform mass univariate linear regressions on “marginal” EEG data

Random-effects contributions are removed from EEG data:

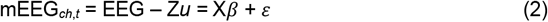

As expressed in (2), what we call marginal EEG (mEEG) can be reconstructed by removing the fitted random values Zu from the data (which is equivalent to adding conditional residuals to trial-wise marginal fitted values). It follows that mEEG can be explained using a (multiple) linear regression model (LM), since we can assume the independence of observations. In the next sections, we will show that the results from (2) are equivalent to fixed-effects results in (1). A single LM is conducted on each channel/timepoint pair. In this way, we obtain a channel-by-timepoint map of the observed *t-*values (*t*-map_OBS_) for each fixed effect. In the proposed example, we obtain a 19×110 *t*-map for the abstract vs. concrete contrast.

### 3. Perform permutation testing and apply threshold-free cluster enhancement (TFCE)

TFCE (Mensen & Khatami, 2013; Smith & Nichols, 2009) is used to assess the significance of *t*-maps_OBS_. First, the design matrix X is permuted thousands of times (e.g., at least 2000 to properly estimate the critical statistics with an alpha level of .05). At each iteration, the permuted X is used for the mass linear regressions as in step 2. Notably, LM are solved using OLS, which is much faster than the RE(ML) method used for LMM, and hence feasible for performing permutation testing in the whole data space (just this simplified example requires 4,180,000 tests). For each effect of interest, TFCE is applied to the corresponding *t*-map_OBS_ and to the *t*-maps obtained from each permutation (*t*-maps_PERM_). The maximum TFCE values from *t*-maps_PERM_ (maxTFCE) are then extracted to build the empirical distribution of maxTFCE values under H_0_, which is used to evaluate the statistical significance of *t*-maps_OBS_. Attention should be paid to two aspects. First, the use of permutation testing is important not only for multiple comparison corrections. It also allows us to overcome the difference between LMM and LM estimations. The fixed-effects *t* values are computed as *β* divided by their standard errors (SEs). As shown in Section 3.3, *β* values are the same between LM and LMM testing. Conversely, since the covariance matrix of fixed-effects coefficients is derived differently in LM as compared to LMM (Bates, Mächler, et al., 2015), SEs differ between LM and LMM, although they are correlated to ∼1. It follows that the *t* values are correlated to ∼1 between LMM and LM, but they differ in value. This aspect is not an issue, since significance is evaluated not based on the absolute *t* value, but based on the empirical distribution of maxTFCE values under H_0_.

## 3. Validation

To validate our analytical strategy, we first assessed its performance characteristics on a univariate simulated dataset (like a behavioral dataset) in terms of power and false positive rate (FPR). Then we applied lmeEEG to a simulated EEG dataset. In the latter case, validity was assessed in terms of equivalence between the lmeEEG results and the results obtained from the highly computationally expensive LMM permutation testing.

The analyses were carried out using MATLAB R2022b on a DELL Precision 7920 Tower, Processor Intel Xeon Gold 5220R (24 cores up to 2.20 GHz), 64 GB RAM, OS Windows 10.

The simulated dataset and MATLAB scripts to perform lmeEEG are available at osf.io/kw87a.

### 3.1. Validation on the simulated univariate datasets

We simulated 2000 datasets from a design with crossed random factors of subjects and items. The design structure is summarized in the following statistical model:

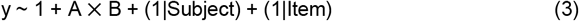

Concerning the fixed effects, A and B were two factors, each including two levels coded as - 0.5 and 0.5 (i.e., effects coding). The A effect had a mean of 0 and a standard deviation (SD) of 1 (i.e., Cohen’s d = 0), that is, a null effect. The B effect had a mean of 0.0695 and SD of 1 (Cohen’s d = 0.07). The interaction between A and B had a mean of 0.05405 and SD of 1 (Cohen’s d = 0.054). Random intercepts for subjects (N = 50) and random intercepts for items (N = 50) had a SD of 0.2, which correspond to a variance partitioning coefficient (VPC) of .2857. Residual errors had a SD of 0.3, which correspond to a VPC of .4286. Each combination of A and B had three repetition within items and subjects. Therefore, each simulated dataset included 30000 observations.

The lmeEEG procedure was applied on each dataset: (1) a mixed model specified as in Equation 3 was conducted; (2) estimated random-effects contributions were removed (see Supplementary Information S1 for the distribution properties of marginal data) and a simple linear regression (i.e. OLS) was conducted on marginal data; (3) permutation testing (without TFCE, since the datasets were univariate) was performed to assess significance of fixed effects. Since the A effect was null, the percentage of significant A effects across simulation gave us an estimation of the FPR (alpha level = .05). Specifically the FPR for A was .045, that is, practically equivalent to the alpha level used, for all the methods. Conversely, the percentage of significant effects for B and the interaction between A and B gave us an estimation of power (alpha level = .05). The power of B was ∼1 compared to the .95 power predicted by the Westfall’s approach (Westfall et al., 2014). The power of the AB interaction was .813 for the permutation test, .828 for the mixed model, and .829 for the OLS on marginal data, compared to the .80 power predicted by the Westfall’s approach (Westfall et al., 2014). To ensure the robustness of the results, this analysis was repeated on simulated datasets with different variance and in a dataset violating the assumption of normality (see Supplementary Information S2 for the analysis description and the results). Overall, lmeEEG showed excellent performance properties.

### 3.2. Simulation of EEG data

Event-related EEG datasets were simulated using the MATLAB-based toolbox SEREEGA (Krol et al., 2018). Epoched data included a P3 potential with different intercepts for subjects (N = 30) and items (N = 10) embedded in noise. Moreover, the P3 of both datasets was modulated differently according to two experimental conditions (i.e., a two-level experimental factor: A vs. B). In detail, for each subject, item, and experimental condition, we simulated 50 epochs of 1100 ms (100 ms of pre-stimulus) at 100 Hz. Each epoch consisted of the sum of the activity of 19 simulated EEG sources spread across the brain and projected onto a standard 19-channel montage. Of the 19 simulated components, 18 were a mixture of white and brown noise (amplitude of each type of noise = 2 μV) with random source locations. The last component was a P3a, whose configuration was taken from the P3a template in SEREEGA. For each grouping factor (i.e., subjects and items), random intercepts were sampled from a normal distribution with 0 μV mean and SD equal to 0.2 times the amplitude of P3a and added to the amplitude of P3a. Concerning the experimental factor, 0.2 μV were added to the amplitude of P3 in the epochs belonging to the experimental condition B.

### 3.3. Validation analyses for simulated EEG data

To validate lmeEEG, we first applied our method as described above. The LMM of Equation 1 was specified as the following Wilkinson-notation formula:

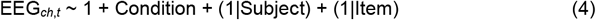

where the fixed-effects part of the model corresponds to 1 for the intercept and Condition is the two-level factor of interest. The random-effects part specifies random intercepts for subjects and items. Next, the mEEG data are reconstructed by adding conditional residuals to the trial-wise marginal fitted values. Finally, mEEG was used to perform steps 2 and 3 described above. In step 3, the design matrix vector was permuted within each subject and item 500 times, and the *ept_mex_TFCE2D* function from the ept_TFCE toolbox (github.com/Mensen/ept_TFCE-matlab; Mensen & Khatami, 2013) was used to perform TFCE.

To validate lmeEEG, step 3 was also performed using LMM on the original EEG dataset. Specifically, the design matrix vector was permuted 500 times (the permuted Condition vector in each permutation was the same between analyses with EEG and mEEG datasets) and the EEG data were explained in each permutation using Equation 3. We limited the number of permutations to 500 (along with simulating a dataset with a small number of channels and time points per epoch) because performing permutation testing with LMM is too computationally expensive (we propose lmeEEG to overcome this limitation) and here we are only interested in demonstrating that the use of our strategy is closely equivalent to performing LMM permutations. The results of both the LMM and LM permutation tests were compared in terms of correlations between *t*-maps_OBS_ and between *t*-maps_PERMS_, as well as in terms of equivalence between the empirical TFCE distribution under H_0_. Finally, lmeEEG performance was assessed using three measures previously used to validate the TFCE approach for EEG (Mensen & Khatami, 2013), namely, sensitivity/power, precision, and the Matthews correlation coefficient (MCC) (Baldi et al., 2000; Matthews, 1975), along with the false positive rate (FPR) (see Supplementary Information S3 for details).

We are assuming that permutation tests with LMM could be considered as the gold standard, given the absence of compelling reasons to cast doubt on it. In fact, the cluster-based correction operates on statistics, which should not depend on the specific statistical method used to generate them (it is a correction for multiple comparisons, not an NHST method). However, given the lack of existing literature on this particular topic, we evaluated the performance of TFCE using LMM, which we present in the Supplementary Information S3.

### 3.4. Validation results of simulated EEG data

As anticipated above, the fitted *β* coefficients were identical when estimated using LMM on EEG data and LM on mEEG data. This aspect was crucial to validate our method. The results showed that both *t*-maps_OBS_ and *t*-maps_PERMS_ had a correlation of ∼1 between the LMM and LM tests (r > 0.99) because, as explained above, the standard errors differed between the two analytical methods, although they were correlated to ∼1. Importantly, however, all the dichotomous decisions based on null hypothesis significance testing (i.e., significant vs. non-significant effects) were the same between the two methods, meaning that there were neither Type 1 nor Type 2 errors. Furthermore, the equivalence of *β* coefficients (i.e., raw effect sizes) between the two methods prevents the possibility of either Type S (sign) or Type M (magnitude) errors (Gelman & Carlin, 2014), ensuring an accurate estimation of experimental effects.

The *p*-value maps obtained from the two procedures were almost identical. Indeed, 98.71% of the *p*-values were identical and 1.1% of the *p*-values differed by one position in the empirical TFCE distributions obtained under H_0_ (the maximum difference was of two positions). This negligible difference in *p-*values, which represents the price of a substantial decrease in computation costs, can nevertheless be reduced by increasing the number of permutations and thus the granularity of the TFCE distributions under H_0_.

Finally, lmeEEG showed a power of .848, a precision of .858, an FPR of .020, and an MCC of .833. Compared to uncorrected, Bonferroni, and FDR corrections, lmeEEG had the best overall performance (see Supplementary Information).

The estimation of LM models for the permutation test (step 3) had a median duration of 0.79 ms (interquartile range IQR = 0.55 ms), while the LMM estimations had a median duration of 256.39 ms (IQR = 49.00 ms). Consequently, our permutation strategy was more than 300 times faster than LMM permutations.

## 4. Application to a real EEG dataset

In this section, we apply lmeEEG to a real EEG dataset collected during a psycholinguistic experiment with a stimuli-within-condition design (Westfall et al., 2014). Fifty-eight students performed a semantic decision task. Participants were asked to decide whether a word presented in the center of the screen denoted an abstract or a concrete concept. The experimental stimuli consisted of 176 Italian words derived from Italian affective norms (Montefinese et al., 2014). During the task, the EEG signal was recorded at a sampling rate of 500 Hz from 58 scalp electrodes mounted on an elastic cap according to the 10–10 International System. Participants’ EEG datasets were preprocessed using an ICA-based pipeline described in Visalli et al. (2021). Clean epochs (from -100 to 1000 ms at 100 Hz) time-locked to word onset were merged across participants. The dimensions of the final EEG dataset were 58 channels × 111 timepoints × 10072 epochs. The LMM was specified as:

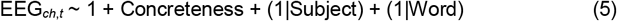

The lmeEEG results are presented in Figure 2. The main finding was a less pronounced N400 event-related potential (ERP) for the abstract compared to concrete words at fronto-central scalp electrodes (Huang & Federmeier, 2015). As in our simulation, the *t*-maps_OBS_ correlation between LM and LMM was >.99.

**Figure 2.**
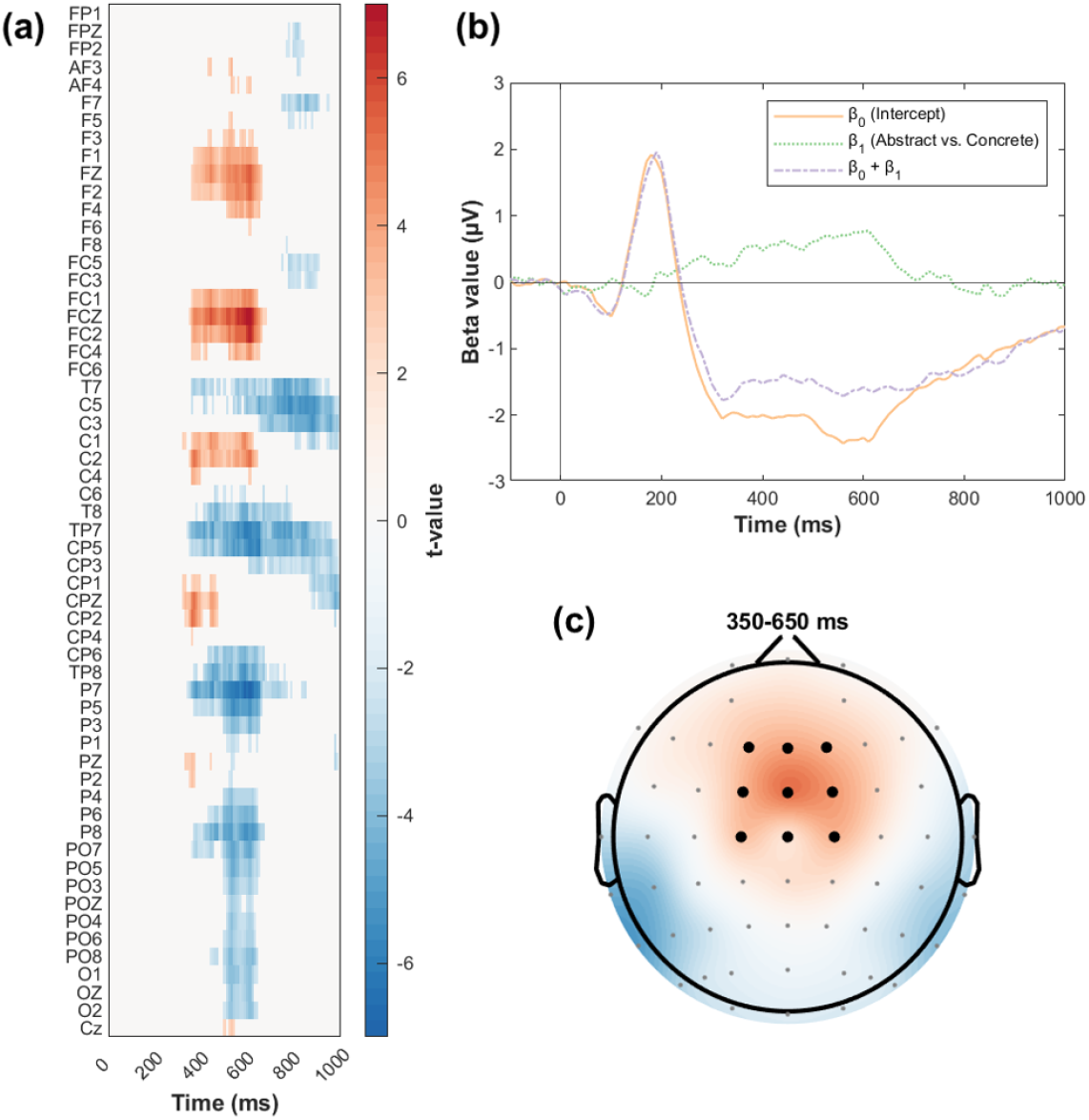
lmeEEG results on the real EEG dataset. (a) Raster diagram showing significant effects elicited by the concreteness predictor. Rectangles in warm and cold colors indicate significantly modulated channel/timepoint pairs. The color bar indicates the *t* values. Gray rectangles indicate electrodes/timepoints for which no significant modulations were observed. (b) Trace-plot depicting the beta values estimated in lmeEEG step 2. Specifically, the intercept (blue line) represents the estimated EEG responses in the “concrete” condition. The *β*_1_ (red line) represents the value to add to the intercept to obtain the estimated EEG responses to abstract words (yellow line). The displayed beta values are averaged across FCz and the eight surrounding electrodes. (c) Topoplot showing the *t* values (same color scale as the raster diagram) averaged in the indicated time window.

## 5. Conclusion

In the present work, we proposed and validated an analytical strategy (lmeEEG) that allows researchers to use mixed models with mass univariate analyses. Essentially, it avoids the unfeasible computational costs that would arise from massive permutation testing with LMM using a simple solution: removing random-effects contributions from EEG data and performing mass univariate LM analysis and permutations on the obtained marginal EEG. Analyses on simulated data showed that the estimated experimental effects and the relative statistical inferences yielded by lmeEEG were equivalent to those obtained by mass univariate analyses with LMM permutations, but almost 250 times faster.

To avoid misinterpretations, the advantages of lmeEEG do not concern accuracy when compared to existing methods for the mass univariate analysis of EEG data (Pernet et al., 2011; Kiebel & Friston, 2004; Ehinger & Dimigen, 2019; Kherad-Pajouh & Renaud, 2015; Frossard, 2019; Frossard & Renaud, 2021). As mentioned in the introduction, they are valuable tools in situations where random coefficient analyses are appropriate. However, they are unable to simultaneously model crossed random effects. Such scenarios require LMM analyses. Unfortunately, LMM analyses are excessively time-consuming for application in mass univariate analysis with permutation testing. Consequently, this challenge has led researchers to either improperly overlook item variability or to use LMM for EEG analyses by performing an (a priori) selection of the data to be tested (thereby avoiding permutations for multiple comparison correction) or some dimensionality reduction procedure (Nielson & Sederberg, 2017). In the present study, we have demonstrated that lmeEEG is a valid, straightforward, and feasible method for conducting LMM across the entire Channel × Timepoint data space. Indeed, the speed advantage offered by lmeEEG overcomes the time-related obstacles associated with employing LMM in a mass analysis approach.

In presenting lmeEEG, we focused on simple ERP studies with one experimental factor and crossed random intercepts for subjects and items. However, our method can be easily applied to a wide variety of experimental studies with more complex fixed structures. Concerning the type of dataset, it can be used for the analysis of even larger EEG data, such as time-frequency data or source-reconstructed ERP, MEG data, or even pupillometry (Montefinese et al., 2018) and eye movement data (Lao et al., 2017). It can also account for designs with “nested” random effects, such as in multi-site neuroimaging studies.

A main drawback of lmeEEG is that it is limited to LMM without random slopes since it requires the permutation of the fixed-effects design matrix (X), but we cannot be sure that random slopes are completely independent of fixed effects. To date, we are not aware of any solution to overcome this issue to apply our approach to random-slope statistical designs. Nonetheless, researchers interested in controlling for inflation of type I error due to the use of random-intercept-only models might apply our method to identify clusters on which to focus the analysis by performing LMM with random slopes. Overall, although not exhaustive, lmeEEG represents a better solution than completely ignoring item variability.

Mixed models are the gold standard for behavioral data analysis in psycholinguistics, where experimental designs always include crossed random variables. Mixed models have several advantages besides modeling crossed random effects, such as, increased power, managing unbalanced datasets or incomplete designs, considering trial-, subject-, and item-related covariates and nested dependencies between data (Baayen et al., 2008). Despite these advantages, mixed models are not yet a common practice in neuroimaging and psychophysiology. Thanks to the possibility offered by the method described in this work, we anticipate that LMM will become increasingly important in neuroscience.

## Ethics approval

Real-data collection was performed in accordance with the ethical standards of the 2013 Declaration of Helsinki for human studies of the World Medical Association. The data collection procedures were approved by the Ethical Committee for the Psychological Research of the University of Padova (protocol number: 2945).

## Data and code availability

Simulation data and codes are available at the Open Science Framework: osf.io/kw87a. Codes and a tutorial for lmeEEG are also available at github.com/antovis86/lmeEEG.

