## Supplementary Information for "lmeEEG: Mass linear mixed-effects modeling of EEG data with crossed random effects"

### S1. Distribution properties of marginal data

In this section, we show that the removal of fitted random effects from the data does not lead to significant alterations in their distribution shape. To this aim, we computed the skewness and kurtosis for the 2000 simulated univariate datasets described in Section 3.1 of the main text. Specifically, we calculated skewness and kurtosis for the datasets both after the removal of random effects (i.e., the marginal data estimated by lmeEEG) and for the same simulated datasets before the inclusion of random effects (i.e., the true marginal data). The results showed that the distributions of the two groups of datasets did not differ significantly either in skewness ($t(1999) = 0.34$, $p = .733$) or in kurtosis ($t(1999) = 0.87$, $p = .382$).

**S2. lmeEEG-validation on additional simulated univariate datasets**

To ensure the robustness of lmeEEG in the face of simulated variance, we replicated the analysis presented in Section 3.1 of the main text on three additional groups of datasets. Each group consisted of 2000 datasets simulated similarly to the main analysis, with the only variation being in the residual errors, which had standard deviations (SD) of 0.6, 1.2, and 2.4, respectively. Table S1 shows the proportion of significant results (positive rate, PR) for the three effects (A, B, and the AB interaction) obtained from each dataset using both conventional linear mixed-effects models (LMM) and linear regression models in marginal data (lmeEEG). Notably, the results remained consistently similar between the two types of analyses, regardless of the simulated variance.

**Table S1 | Positive rate (PR) for LMM and lmeEEG analyses on simulated datasets with different variance.**

| SD of residual errors | Effect | LMM PR | lmeEEG PR |
|---|---|---|---|
| 0.6 | A | .053 | .053 |
| | B | 1 | 1 |
| | AB | .678 | .679 |
| 1.2 | A | .053 | .053 |
| | B | .989 | .989 |
| | AB | .371 | .371 |
| 2.4 | A | .057 | .057 |
| | B | .665 | .666 |
| | AB | .161 | .161 |

Finally, to confirm the robustness of lmeEEG when faced with violations of normality assumptions, we conducted the same analysis on a group of datasets with log-normal distribution (an exponential transformation was applied to the datasets described in Section 3.1). Also in this case, PR remained nearly identical between LMM and lmeEEG. Specifically, the A effect had a PR of .053 for both analyses, the B effect had a PR of ~1 for both analyses, and the AB interaction had a PR of .651 for LMM and .652 for lmeEEG.

**S3. Assessment of threshold-free cluster-enhancement (TFCE) with LMM.**

To evaluate the performance of the TFCE[1] approach using LMM, we compared TFCE with other non-parametric corrections, namely, Bonferroni and False Discovery Rate (FDR)[2].

To summarize, the following LMM was conducted on each channel/timepoint combination of the simulated EEG dataset:

$$EEG_{ch,t} \sim 1 + \text{Condition} + (1|\text{Subject}) + (1|\text{Item})$$

In this way, we obtained a channel-by-timepoint map of the observed *t* values ($t$-map$_{OBS}$) for the Condition effect, as well as the associated *p*-values ($p$-map$_{OBS}$). The fixed-effects design matrix (X) was then permuted within subjects and items 500 times. At each iteration, the permuted X was used to carry out the LMM specified above, and a *t*-map ($t$-map$_{PERM}$) was obtained for each permuted dataset. Finally, TFCE was applied on the $t$-map$_{OBS}$ and $t$-maps$_{PERM}$. The maximum TFCE values of each $t$-map$_{PERM}$ (maxTFCE) were then extracted to build the empirical distribution of the maxTFCE values under $H_0$, which was used to evaluate the statistical significance of the $t$-map$_{OBS}$.

The TFCE results were compared to those obtained from Bonferroni and FDR corrections applied to the $p$-map$_{OBS}$ in terms of the measures previously used to validate the TFCE approach for EEG[1], namely, sensitivity/power, precision, and the Matthews correlation coefficient (MCC)[3,4], along with the false positive rate (FPR). Specifically, power was computed as the ratio between the number of true positives (TP) found and the actual number of positive channel/timepoint pairs (TP and false negatives, FN). The precision was computed as the ratio between the number of TP and the total number of positives found (TP and false positive, FP). The FPR was computed as the ratio between the number of FP and the number of negative channel/timepoint pairs (FP and true negative, TN). The MCC was defined as follows:

$$MCC = \frac{TP \times TN - FP \times FN}{\sqrt{(TP+FP)(TP+FN)(TN+FP)(TN+FN)}}$$

All these measures were calculated for the statistical threshold criteria of $p = 0.05$.

The computation of these measures requires the definition of the relevant signal (i.e., the channel/timepoint pairs where the Condition-dependent difference in the P3a signal was present). Since the P3a were simulated at the dipole source and then projected onto the scalp, we simulated the P3a of the two levels of the Condition factor without noise and then computed the difference on the scalp. The relevant signal should be any channel/timepoint pair where the difference is not zero. However, since the simulation occurred at the dipole source, virtually any channel/timepoint datapoint presented a nonzero effect. Therefore, following Mensen and Khatami[1], the relevant signal was defined as the channel/timepoint pairs whose absolute Condition effect was greater than 12.5% of the maximal effect (contour region in Figure S1).

Performance metrics are shown in Table S2. Compared to uncorrected data, Bonferroni, and FDR corrections, TFCE had the best overall performance. Table S2 also presents performance metrics for lmeEEG.

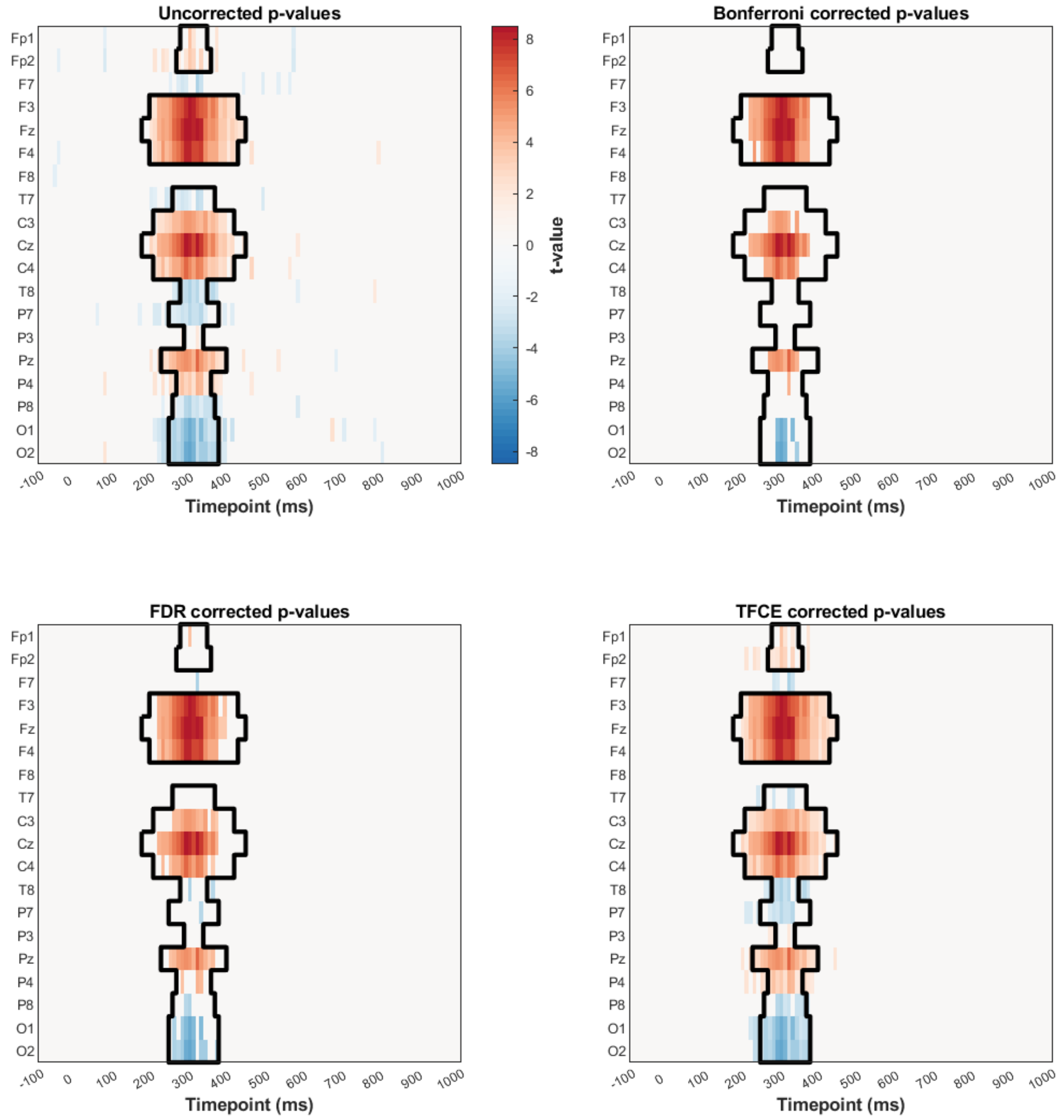


**Figure S1|** Raster diagrams show significant effects in our simulated experiment according to different correction procedures. Rectangles in warm and cold colors indicate channel/timepoint pairs that were positively and negatively significantly modulated, respectively. The color bar indicates the *t* values. Gray rectangles indicate electrodes/timepoints for which no significant modulations were observed. Significance was set at the *p*-value threshold of 0.05 adjusted according to the indicated correction. The contour regions indicate the relevant signal (channel/timepoints pair with a true effect greater than 12.5% of the maximum effect).

**Table S2 | Performance metrics**

| Corrections | Power | Precision | FPR | MCC |
|---|---|---|---|---|
| **Uncorrected** | 0.8755 | 0.7352 | 0.0441 | 0.7722 |
| **Bonferroni** | 0.3670 | 1 | 0 | 0.5827 |
| **FDR** | 0.4864 | 0.9843 | 0.0011 | 0.6671 |
| **TFCE** | 0.8483 | 0.8583 | 0.0196 | 0.8328 |
| **lmeEEG** | 0.8483 | 0.8583 | 0.0196 | 0.8328 |

*Notes:* MCC: Matthews correlation coefficient; FPR: false positive rate; FDR: false discovery rate; TFCE: threshold-free cluster-enhancement; lmeEEG: mass univariate linear mixed-effects modeling of EEG data.
